# Task-Relevant Cognitive Load Modulates Fast and Slow Processes Underlying Motor Adaptation

**DOI:** 10.64898/2026.09.17.752503

**Authors:** Vyaas Ramasubramanian, Somesh Nana Shingane, Aditya Murthy

## Abstract

How cognitive control influences the correction of motor errors is an important question. In this study, we examined how the slow and fast processes that constitute motor adaptation are modulated by cognitive load. To address this question, we used a novel approach that differed from traditional dual-task experiments that have assessed the role of fast processes by decreasing attention to the motor adaptation task. Here, cognitive load was manipulated using a decision-making task with easy and hard difficulty levels, while simultaneously adapting to a perturbation and maintaining attention on the task. Our results showed that the hard decision-making task increased cognitive load relative to the easy task, resulting in attenuated steady-state learning. The relative contributions of the fast and slow components were assessed using a dual state-space model, which showed that the former was larger under the high-load condition, whereas the latter was larger under the easy-load condition, consistent with a competition model in which a shared error signal is partitioned between the two processes. We also showed that the effect of cognitive load on fast processes was not attributable to increased reaction times or task performance. Taken together, these results support the notion that faster explicit motor adaptation processes derive from cognitive processes involved in action selection.

**NEW AND NOTEWORTHY:** Prior dual-task studies examining the effect of cognitive load on motor learning have shown that the faster explicit process is negatively affected by higher load. We have shown that when attention and cognitive load are not divided between two tasks but instead focused on the motor task, the effect is reversed, and the expression of the faster explicit process is boosted. Interestingly, the greater expression of the faster explicit process did not lead to greater adaptation in the learning phase, consistent with the notion that the slower implicit and faster explicit processes are also yoked to a common error.

## INTRODUCTION

Motor adaptation reflects the process by which the motor system modifies its output to compensate for changes in the body or environment. This adaptive behavior is thought to emerge from the contributions of fast and slow learning processes (Smith et al., 2006). The fast process learns rapidly from error but exhibits poor retention, whereas the slow process adapts more gradually and retains learning over longer periods (Smith et al., 2006). In addition to representing distinct computations, these two processes are also thought to be driven by two distinct errors, suggesting the fast and slow processes reduce task errors and sensory prediction errors, respectively, in an independent manner (Mazzoni and Krakauer, 2006; Taylor and Ivry, 2011; Wong and Shelhamer, 2012). In contrast, recent studies have suggested a competitive interaction between these two processes, supporting a shared performance error (Albert et al., 2022; Neville & Cressman, 2018; Saijo & Gomi, 2010).

The two-state framework has also been interpreted in terms of explicit and implicit learning mechanisms. The fast process is thought to reflect strategic, cognitively driven adjustments in behavior and is often associated with an explicit mode of learning. For example, task errors and the increase in the use of cognitive strategies, resulting in longer preparation time, awareness of the perturbation, and explicit instructions, are known to influence the fast process (Fernandez-Ruiz et al., 2011; Haith et al., 2015; Leow et al., 2020; Neville & Cressman, 2018; Sadaphal et al., 2022). In contrast, the slow process is believed to represent a more automatic recalibration of the sensorimotor system, and is thought to be driven by primarily driven by sensory prediction errors which reflects an implicit or automatic mode of learning (McDougle et al., 2015; Taylor & Ivry, 2011) since it persists even when compensation is discouraged or the perturbation is task-irrelevant (Kim et al., 2018; Mazzoni & Krakauer, 2006; Morehead et al., 2017; Shingane et al., 2025).

In traditional dual-task paradigms, attention is divided between tasks (McIsaac et al., 2015; Taylor & Thoroughman, 2007; Woollacott & Shumway-Cook, 2002), while in natural behavior, cognitive demands are often embedded within the motor task itself rather than imposed externally. In natural settings, however, motor adaptation rarely occurs in isolation. Rather, it occurs alongside concurrent cognitive demands and competing task requirements. To capture this complexity, dual-task paradigms have been widely used, in which participants perform a secondary cognitive task during adaptation. These studies have shown that increasing cognitive load largely spares implicit learning, while selectively impairing explicit learning, consistent with the notion that explicit processes depend on limited cognitive resources. In contrast, implicit processes operate more automatically and are immune to such cognitive manipulations (Keisler & Shadmehr, 2010; Zhang et al., 2025).

To address whether fast explicit and slow implicit learning can also be modulated by task-relevant cognitive load and test whether these processes compete for a common error during learning, the present study introduces cognitive load intrinsically within the motor task by requiring participants to make target-selection decisions during movement execution. This approach allowed us to manipulate cognitive load while maintaining attention within the same action space. Using this approach, it was observed that although the overall adaptation was comparable across the two cognitive loads, the relative contributions of faster explicit and slower implicit processes differed. Specifically, higher cognitive load increased the contribution of explicit process while reducing the slower implicit adaptation, whereas lower load showed the opposite pattern. Thus, when cognitive demands are integrated within the motor task, they may actively engage strategic processes. Furthermore, these results support the notion that explicit and implicit learning compete for a shared error signal and show how embedded cognitive loads influence motor adaptation.

## METHODS

### Participants

A total of 40 naive participants (29 males, 11 females, ages 18-34) with normal or corrected-to-normal vision and no history of motor impairments participated in the experimental study, which was approved by the Institute Ethics Committee at IISc. All participants were right-handed, as verified by the Edinburgh Handedness Inventory (Oldfield, 1971). Each experimental condition included 20 participants, and participants were randomly assigned to one of the conditions.

### Experimental Apparatus

Stimulus presentation and data collection were performed using NIMH MonkeyLogic (Hwang et al., 2019) in MATLAB (MathWorks Inc.). Reaching movements were performed on a 21.5-inch XP Pen Artist Display 22R Pro tablet (Hanvon Ugee Group, Shenzhen, China) with a resolution of 1920 × 1080 pixels at 60 Hz using a pressure-sensitive, battery-free stylus. The positional and event data were continuously collected by MonkeyLogic. The participants observed the task through a monitor placed on a desk that obscured the XP Pen and was of the same dimensions (Figure 1A; top).

**Figure 1:**
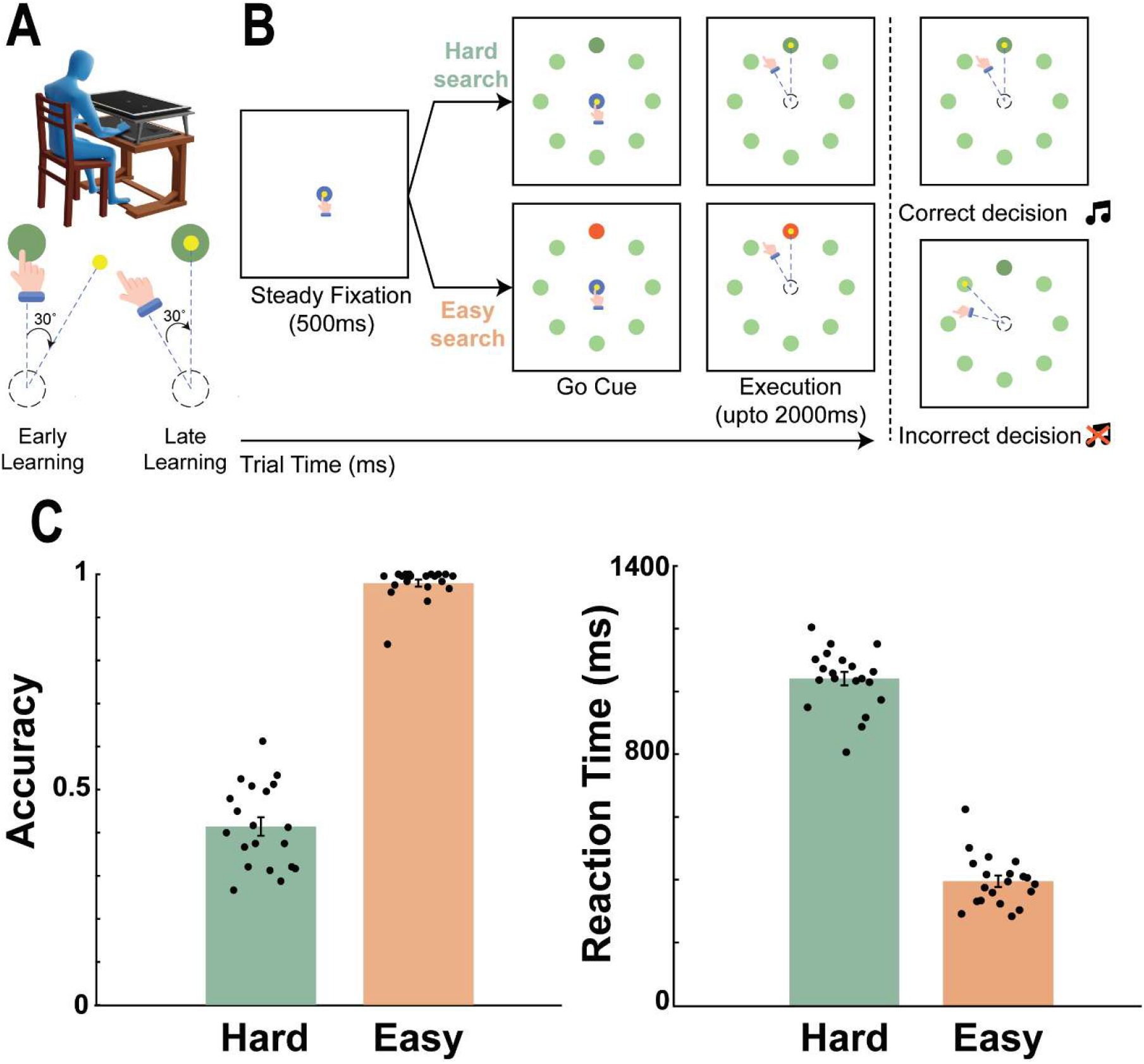
**(A) Experimental setup and rotation perturbation design:** Participants performed a reaching task to a target by making a planar movement with a digitized stylus in their right hand, which is reflected by the cursor movement on the screen (Top). During the perturbation, the participant experienced a 30-degree clockwise rotation offset between the hand (hand icon) and the cursor (yellow) movement while reaching to the desired target/distractor (green) from the starting position (blue dotted circle) creating movement error in early learning session (bottom left) which the subject learns to compensate for by the end of the perturbation period (bottom right). Note that for both tasks, the rotation perturbation between hand and cursor was not presented during the baseline and washout period; the cursor feedback was veridical to the hand movement. **(B)Combining visual search task with visuomotor perturbation:** Participants performed a visual search task by reaching to a target as embedded among distractors, where the search for the target could either be an Easy search task (red target among green distractors) or a Hard search task (darker green target among green distractors). During the perturbation period, the participants experienced a 30-degree clockwise offset between hand and cursor movements. For both tasks, each trial would start with steady fixation (500 ms) of the cursor (hand) at the start position. Following successful fixation, participants are immediately presented with the stimulus (target and distractors), marking a go-cue to execute the movement to the selected target/distractor location within the time limit of 2000 ms (left panel). Participants would hear a tone upon a correct selection (right panel, top) and no tone upon an incorrect selection (right panel, bottom). Note that the dotted lines following the cursor and hand, and the dotted circle at the start position, are for illustrative purposes; participants did not observe such feedback during the task. The fixation circle has been illustrated in blue here for ease of visualization. **(C)Accuracy and Reaction time performance:** The accuracy (left) and reaction time (right) in the Easy (tangerine) and Hard (green) search task indicate that cognitive load decreased accuracy but increased reaction time during the perturbation period. The height of the bars reflects the mean, while the error bars reflect the SEM. For both plots, each dot represents data from an individual subject.

### Task Design

Participants performed the reaching task by dragging the stylus on the tablet surface. In parallel, a screen of the same dimensions was mounted, which displayed the stylus position as a cursor. This setup occluded the participant’s hand during reaching, allowing us to introduce a rotation perturbation by creating a spatial mismatch between the hand and the cursor feedback (Figure 1A; top). For the baseline trials, the hand position was mapped onto a yellow cursor, and continuous feedback was provided throughout the movement. The rotation perturbation was introduced during the task’s perturbation period, which participants were unaware of (Figure 1A; bottom). The task involved a visual search in which participants were instructed to make reaching movements toward the odd target among 8 possible locations. The degree of task difficulty was manipulated across two conditions: easy and hard search. In both conditions, the seven distractors were green (CIE: (x, y = 0.28, 0.59, 15.0 cd/m^2^)) and the odd target differed in colour (Figure 1B). In the easy condition, the target was red (x, y = 0.66, 0.28, 4.70 cd/m^2^), making it highly salient, while in the hard condition, the odd target was of a darker shade of green (CIE: (x, y = 0.29, 0.56, 7.7 cd/m^2^)) and thus, harder to identify (Figure 1B). The position of the odd target was pseudo-randomized, appearing at any of the 8 locations during a cycle of 8 trials. Participants in the hard group were instructed to make movements toward a random target if they were unable to find the odd target in a trial, to avoid skipped trials (11.61% ± 0.068). Participants were randomly assigned to either group.

### Experiment Structure

Before the experiment, all participants completed 40 practice trials moving to a single target to become accustomed to the required movements. The experiment consisted of 3 periods: a baseline period, a perturbation period, and a washout period. The experiment consisted of 45 cycles, with 8 trials per target location, and the correct order was randomized, for a total of 360 trials. The baseline consisted of 5 cycles. Following the baseline, a 30° clockwise perturbation was introduced using the cursor. Participants were not informed of the upcoming perturbation.

The perturbation period lasted for 30 cycles. Finally, there was a 10-cycle washout period during which the perturbation was removed (Figure 2A).

**Figure 2:**
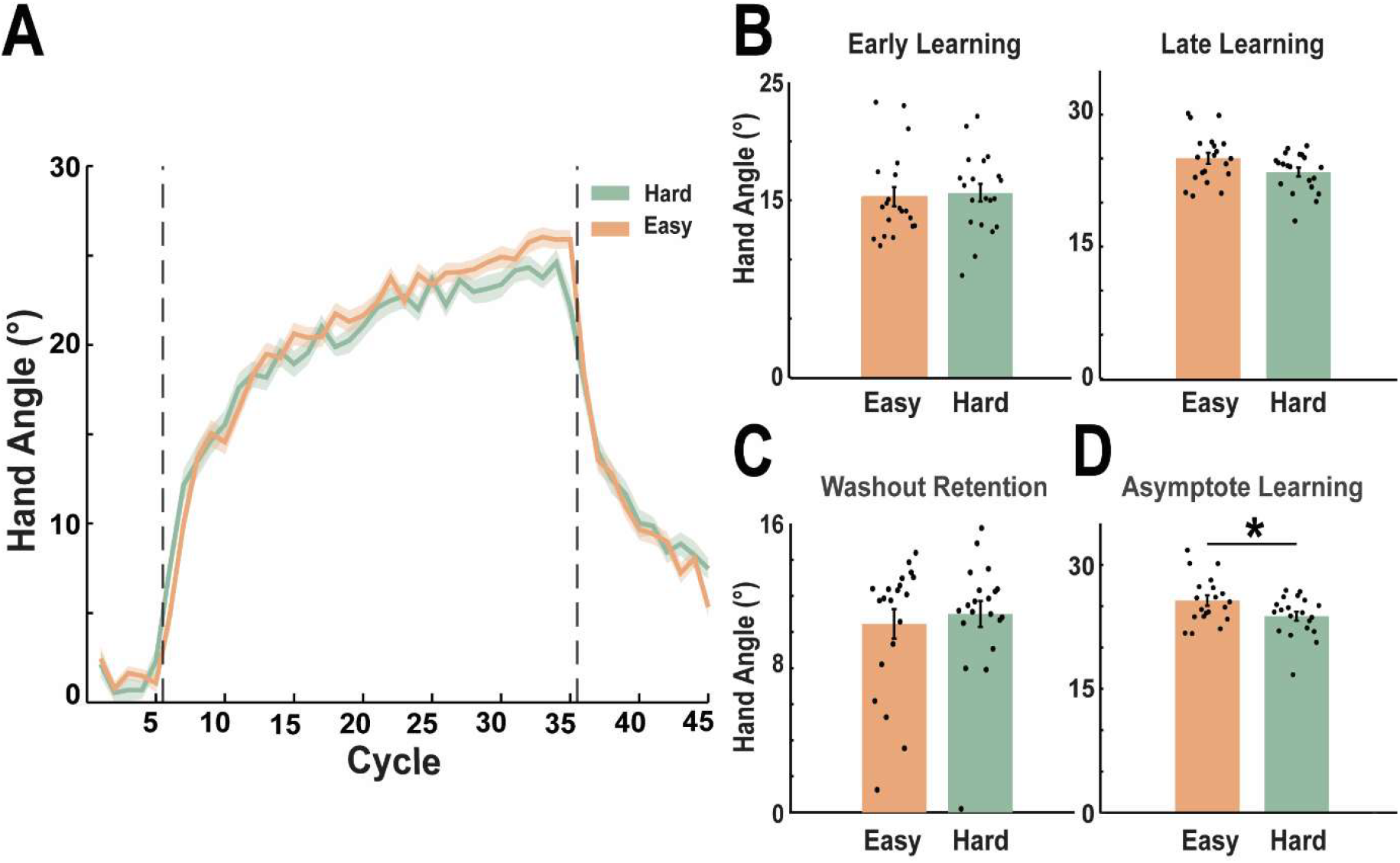
Learning curve and performance for visuomotor rotation perturbation for easy (tangerine) and hard (green) search conditions. **(A)** Both the easy and hard conditions show similar learning but reach different asymptotic levels. The dashed vertical lines indicate the start and end of the perturbation period. The dark coloured solid lines indicate the mean, and the shaded region indicates the standard error. **(B) (C) and (D)** Condition-averaged hand angle at early and late learning during the perturbation period, at the early washout period (first 10 cycles), and asymptotic learning (perturbation period: last 5 cycles), respectively. Learning at the early, late, and washout stages shows no difference, whereas the asymptote stage shows greater learning in the Easy condition than in the Hard search condition. The height of the bars reflects the mean, while the error bars reflect the SEM, and each dot represents individual participant data.

### Trial Design

A white square at the centre indicated the fixation point for every trial. Participants had to place the cursor on the fixation square and, upon holding it at this position for 500ms, the 8 targets appeared at 8 cm from the centre, evenly spaced at 45 ° apart. Participants then had 2000ms to identify the odd target and make the movement. Participants had to move the cursor to the chosen target and hold it for 500ms (Figure 1B). If the correct target was chosen and the cursor was held in the target window for 500ms, an auditory tone was presented indicating the correct choice (Figure 1B; right). There was no feedback for choosing an incorrect target. Upon completion of the trial, the screen would go blank for 2000ms, meanwhile participants would bring their hand back towards the centre of the screen.

### Data Analysis

All data acquisition and analysis were conducted using MATLAB (MathWorks Inc.) and associated software, including NIMH MonkeyLogic (Hwang et al., 2019). Positional data were acquired via MonkeyLogic sampling at 1 kHz. The positional data from MonkeyLogic was further low-pass filtered using a fourth-order Butterworth filter with a 20Hz cutoff. The filtered data were differentiated successively to obtain kinematic parameters such as velocity, acceleration, and hand angle.

#### Hand Angle

The hand angle was calculated at the point of peak velocity. The angle between the vectors from the centre to the target and from the centre to the point of peak velocity gave the reach angle. Anticlockwise reaches countering the clockwise perturbation were positive reach angles, while clockwise reaches were considered negative reach angles. Hand angles exceeding 45° were excluded to avoid movements that reflect ambiguous decisions.

#### State Space modelling

The dual state space model as proposed by Smith et.al (Smith et al., 2006) was used to model the slow and fast processes, which are thought to serve as proxies for the implicit and explicit processes, respectively. (McDougle et al., 2015)

The model is formulated as:

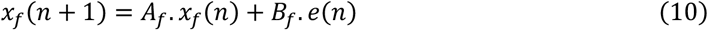

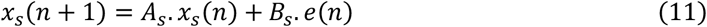

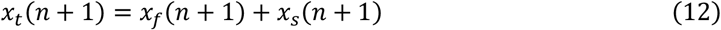

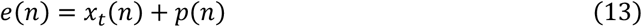

With the constraints,

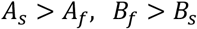

where,

n is the cycle number

*x*_*t*_(*n*) is the hand angle on cycle n

*x*_*f*_(*n*) contribution to the fast process for cycle n

*x*_*s*_(*n*) contribution to the slow process at cycle n

*A*_*f*_ and *A*_*s*_ are the retention factors with bounds [0,1]

*B*_*f*_ and *B*_*s*_ are the learning rate/error sensitivity parameters with bounds [0, 1]

*e*(*n*) is the error experienced due to perturbation p(n) at cycle n

The parameters were estimated for both the individual and the population average for both conditions. The fits were performed using the fmincon function in MATLAB, with squared error as the objective function. The contributions of implicit and explicit learning were further quantified using peak magnitude and area under the curve (AUC) from characteristic curves generated by the retention and learning-rate parameters.

The model was fit to individual participants to assess whether these trends were observed in them. The peak of slow and fast learning, as well as the area under the curves of both processes during the perturbation period, were also estimated. There were 2 outlier participants in the easy condition, who were excluded from this analysis. For these two participants, the learning curve was accounted for entirely by the fast process, with the slow component plateauing at approximately 5 ° in contrast to the remaining participant’s peak slow learning (hard: 21.04 ° ± 3.25 °, easy: 23.09 ° ± 3.31 °).

To examine how the outcome of a decision influenced subsequent adaptation in the hard condition, trials were first classified as correct or incorrect based on whether the participant selected the correct target. For each trial, the next trial in which the participant reached the same target location was then identified. The movement angle in the subsequent trial was extracted and compared with that from the original trial. For example, if a participant selected the upper-left target on trial 15, the movement angle on trial 15 was paired with the movement angle from the next trial in which the upper-left target was selected (e.g., trial 22). The relationship between the movement angle on the initial trial (*a*_t_(*q*)) and the angle on the subsequent trial to the same target (*a*_*t*_(*q* + 1)) was modelled as

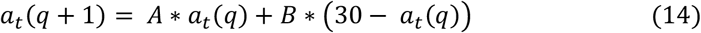

where (30 °) corresponds to the imposed visuomotor rotation. In this formulation, (A) quantifies retention and (B) quantifies the learning rate/error sensitivity. The parameters (A) and (B) were estimated separately for trials following correct and incorrect decisions using least-squares regression. Comparisons of the resulting parameter estimates were used to assess whether the decision outcome influenced retention and error-based updating. The perturbation period was investigated in this analysis.

#### Statistical testing

ANOVAs were performed to test main and interaction effects. A Shapiro-Wilk test was used to test for normality. If normality was satisfied, a t-test was used; otherwise, a Wilcoxon rank-sum test was used to check for significance. When reporting results, normal distributions are reported as the mean and standard deviation (mean ± SD), and non-normal distributions are reported as the median, including the interquartile range (IQR).

#### Accuracy

This was defined as the proportion of times the correct target for reaching was chosen.

## RESULTS

Participants performed a visuomotor reaching movement coupled with a visual search task of easy or hard difficulty. During the perturbation period, participants experienced a 30° clockwise perturbation while reaching towards the chosen target location. Performance in the easy condition was nearly perfect, and participants performed significantly better than in the hard condition (hard: 0.41 ± 0.1, easy: 0.996 (0.97-1), W = 210, p < 0.0001, r = 0.86; Figure 1C; right). The performance under the hard condition was also higher than 0.125, indicating an effort to perform the task rather than to guess. However, for a few participants, baseline performance was near chance, which subsequently increased across the various phases of the task. Reaction times were also measured to assess speed-accuracy trade-offs. Reaction times were significantly higher in the hard condition (hard: 1041 ± 96 ms, easy: 395 ± 81 ms, t = 22.9381, p < 0.0001, Cohen’s d = 7.11; Figure 1C; left). This simultaneous increase in reaction time and decrease in search accuracy in the hard condition is indicative of a modulation of cognitive load rather than a speed-accuracy trade-off.

As expected, participants successfully adapted to the visuomotor perturbation across all task conditions, as reflected in their hand angle during the perturbation period (Figure 2A). To investigate differences in adaptation between the two conditions, the learning period was divided into 3 stages: early, middle, and late (30 cycles total; 10 cycles each). No significant effect of task condition across the different stages of the perturbation period was found (p = 0.46, ANOVA, F = 0.5589; Figure 2B). However, the test revealed a possible interaction between condition and stage of learning (p = 0.089, ANOVA, F = 2.49). To assess this further, the last 5 cycles of the perturbation period were treated as the learning asymptote (Figure 2D). The comparison revealed an effect of task condition on learning at the asymptote, with larger steady-state learning in the easy condition (hard condition: 23.79 ± 2.45, easy condition: 25.68 ± 2.82, t = 2.26, p = 0.03, Cohen’s d = 0.7). Taken together, these results indicate that while learning largely remained similar between the two conditions, the effect of cognitive load on learning was expressed at the steady state and adaptation was reduced with higher cognitive load.

To further investigate whether such steady-state differences in cognitive load also altered the expression of slow and fast processes, a dual state-space model was used to estimate the fast and slow processes for the group means and individual participants. The group fit of the dual state space model revealed a higher expression of the fast process in the hard condition relative to the easy condition (Hard Condition: A_f_ = 0.41, B_f_ = 0.28, peak = 8.77°, AUC = 142.18; Easy Condition: A_f_ = 0.37, B_f_ = 0.24, peak = 7.40°, AUC = 107.61) and vice versa for the slow process (Hard Condition: A_s_ = 0.98, B_s_ = 0.09, peak = 21.71°, AUC = 436.54; Easy Condition: A_s_ = 0.98, B_s_ = 0.12, peak = 23.08°, AUC = 492.11) (Figure 3A, 3B). To assess the robustness of this result, the washout and late learning stages (the last 10 cycles) were removed, and the data were refitted; the trends remained similar. (Washout Removed: Hard Condition: A_f_ = 0.37, B_f_ = 0.29, fast peak = 8.69°, fast AUC = 133.78, A_s_ = 0.97, B_s_ = 0.11 slow peak = 20.94°, slow AUC = 451.93; Easy Condition: A_f_ = 0.52, B_f_ = 0.23, fast peak = 8.26°, fast AUC = 130.56, A_s_ = 0.99, B_s_ = 0.10, slow peak = 23.53°, slow AUC = 475.56; Late Learning Removed: Hard Condition: A_f_ = 0.38, B_f_ = 0.29, fast peak = 8.76°, fast AUC = 104.81, A_s_ = 0.97, B_s_ = 0.11, slow peak = 19.20°, slow AUC = 247.09; Easy Condition: A_f_ = 0.43, B_f_ = 0.22, fast peak = 7.08°, fast AUC = 82.24, A_s_ = 0.97, B_s_ = 0.12, slow peak = 20.86°, slow AUC = 274.78) The parameters obtained from the model fitting are shown in Table 1.

**Table 1.**
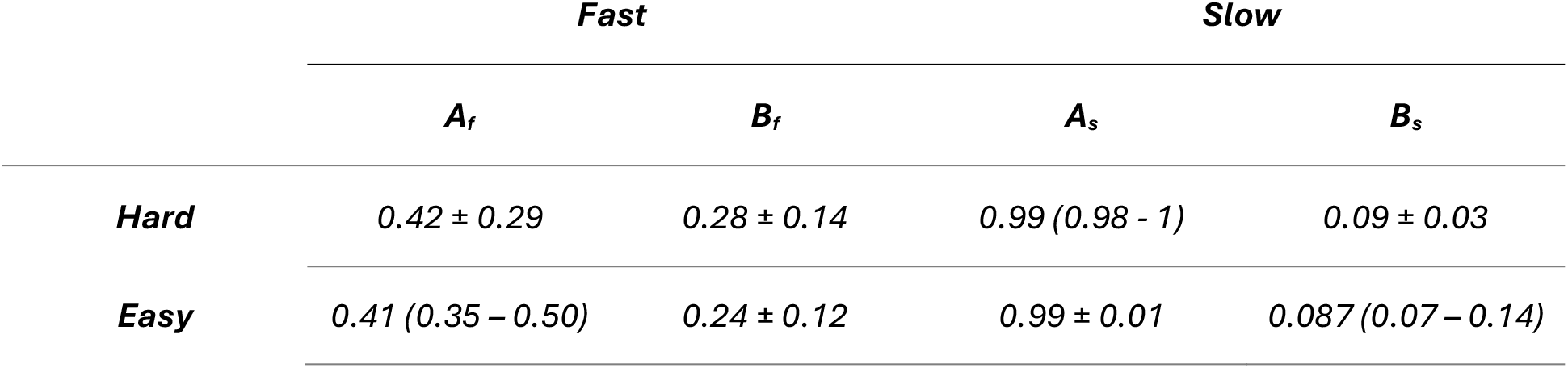
The parameters obtained from the SSM for both the easy and hard conditions.

**Figure 3:**
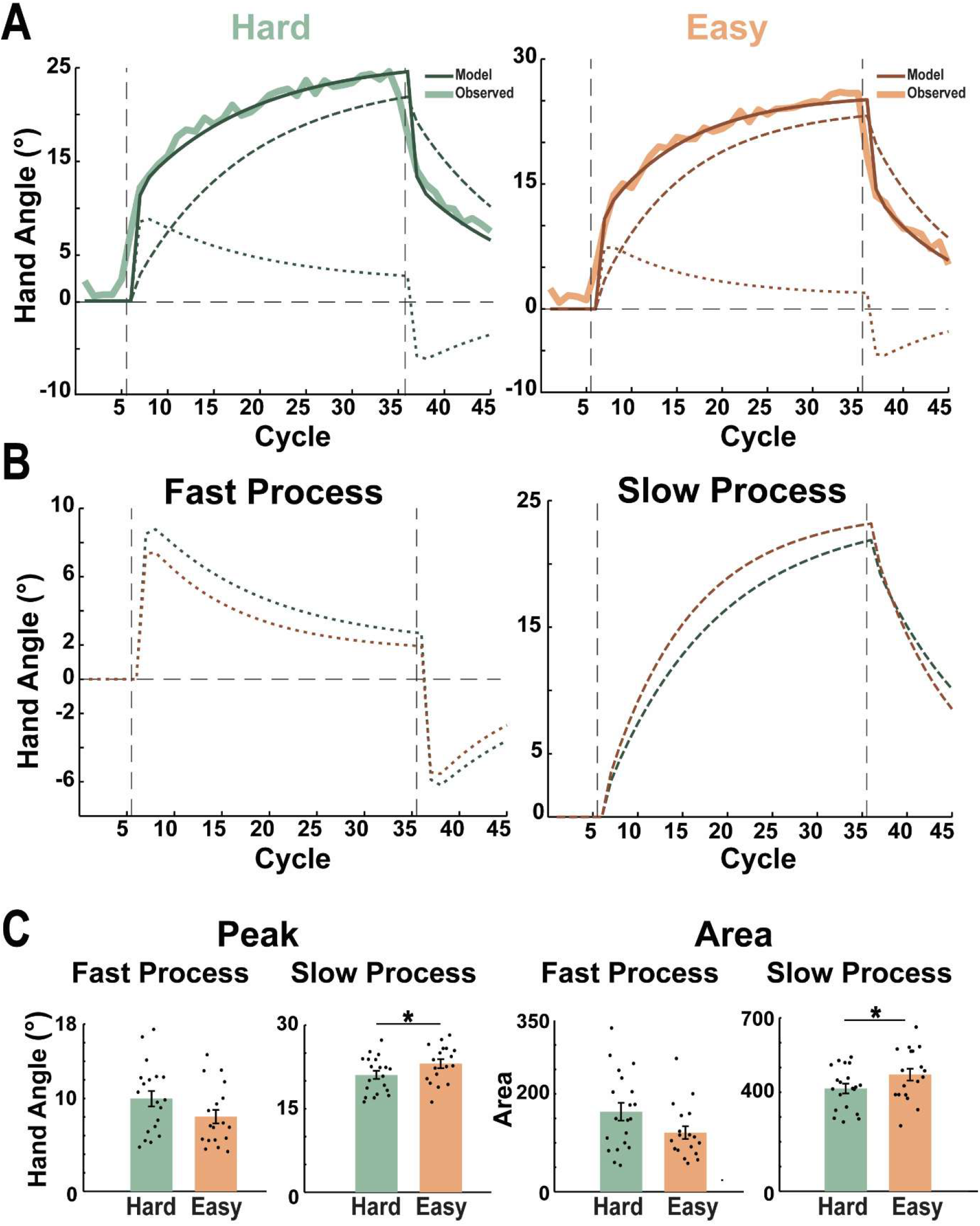
Dual state-space model fitting and processes comparison for Easy (tangerine) and Hard (green) search conditions. **(A)** The model-fitting results for the group averages in the Hard (left) and Easy (right) conditions. The obtained overall learning is indicated by solid dark-colored lines, and the individual slow and fast processes are indicated by dashed and dotted lines, respectively. **(B)** Comparing the dynamics of fast(left) and slow(right) processes for easy and hard search task conditions obtained from dual state space model fitting throughout different periods of the task. In **(A) and (B)**, the vertical dashed lines (black) indicate the duration of the perturbation period, i.e., following the baseline period and preceding the washout period, and the horizontal dashed line indicates zero-degree hand angle reference. **(C)** For the perturbation period, condition-averaged data of peak hand angle value (left) and area under the curve (right) for both the slow and fast processes when individual participants’ data were fit to the two-state model for respective task (easy/hard search) conditions. The height of the bars reflects the mean, while the error bars reflect the SEM, and each dot represents individual participant data.

Individual participants’ behavioral data were also fitted to assess the significance of the results. The peak of learning and the area under the curve were examined for both the fast and slow processes (see Methods). For the fast process, both peak (hard: 9.9987° ± 3.66°, easy: 6.98° (5.53° – 10.03°), W = 445, p = 0.055, r = 0.26) and area under the curve (hard: 163.35 ± 78.77, easy: 108.17(84.78 – 155.75), W = 445, p = 0.055, r = 0.26) were marginally significant (Figure 3C). For the slow process, both peak (hard: 21.04° ± 3.25°, easy: 23.09° ± 3.31°, t = 1.93, p = 0.03, d = 0.61) and area under the curve (hard: 414.49 ± 84.59, easy: 471.17 ± 104.68, t = 1.84, p = 0.037, d =0.59) were significant (Figure 3C). This indicates that cognitive load can modulate the individual processes that comprise motor adaptation, with the fast process being more strongly expressed in the hard condition and the slow process in the easy condition. Since learning across conditions was comparable, we infer that these changes in both slow and fast expression are consistent with the notion that these processes are yoked to a common error signal. A negative correlation between slow and fast area-under-the-curve learning across subjects (Linear Regression, slope = −0.96, intercept = 578.12, r^2^ = 0.468, p < 0.001) supported this notion.

To assess the potential effects of reaction time on the expression of the fast process, the peak of fast learning was correlated with the reaction time observed in the cycle in which it occurred. The correlations revealed that the reaction time was not a predictor of explicit learning. The correlations were not significant for the hard condition (r^2^ = 0.019, p = 0.559), whereas they were significant for the easy condition but did not explain much variance (r^2^ = 0.295, p = 0.02). We also tested whether differences in learning might arise from performance on the decision-making task, especially for the hard search task, as the easy task had an accuracy of 99%. For this, learning following correct and incorrect trials was estimated using equation 14 (see methods) after model fitting. We found that for the hard search task, there were no differences in the area under the curve (correct: 594.67 ± 61.08, incorrect: 585.92 ± 60.56, t = 0.46, p = 0.74) and maximal learning (correct: 21.4611 ± 2.23, incorrect: 20.99 ± 2.1, t = 1.12, p = 0.28). Thus, the effects of increased explicit learning expression were not due to an interaction between the decision-making task and the adaptation processes resulting from increased action errors.

## DISCUSSION

The present study examined how task-relevant cognitive load influences motor adaptation by embedding cognitive demands directly into the motor task, rather than using traditional dual-task paradigms, in which adaptation occurs concurrently with an unrelated cognitive task. This approach allowed cognitive load to be manipulated while maintaining attention within a common behavioral context. Although overall adaptation trajectories were largely comparable between conditions, increasing task difficulty altered the composition of learning. Specifically, the high-load condition was associated with a greater contribution of the fast-learning process, commonly interpreted as reflecting explicit adaptation. In contrast, the low-load condition showed relatively greater expression of the slow learning process associated with implicit adaptation (McDougle et al., 2015). Interestingly, participants in the hard condition reached a lower asymptotic performance level, suggesting that the increased explicit learning did not lead to greater total adaptation. These findings suggest that when cognitive demands are integrated with the motor task, explicit learning is facilitated, thereby shifting the balance between explicit and implicit adaptation, which in turn constrains the steady-state extent of motor adaptation.

### Modulation of fast and slow learning by load

A central finding of this study is that task-relevant cognitive load enhanced the expression of the explicit process. This result contrasts with the prevailing conclusions from dual-task motor adaptation studies. Previous work has demonstrated that a cognitively demanding secondary task reduces explicit adaptation, leaving implicit adaptation relatively unaffected (Keisler & Shadmehr, 2010; Taylor & Thoroughman, 2007; Zhang et al., 2025). This finding has been interpreted as evidence that explicit adaptation depends on limited cognitive resources that must be shared with other concurrent tasks. In such paradigms, however, the cognitive task is typically unrelated to the motor adaptation process, requiring the participant to divide attention across competing goals. Under these circumstances, explicit strategy formation may be impaired because the cognitive resources required to evaluate errors, formulate corrective actions, and update movement plans are consumed by the secondary task.

The present task differs from previous work in that the cognitive demands were integrated with action selection since the visual search process determined the movement goal itself, thereby coupling perceptual decision-making and motor planning into a unified behavioural context. Rather than competing with adaptation for cognitive resources, the decision-making process may have increased engagement with movement planning and outcome evaluation. Explicit adaptation has been linked to conscious strategy generation, action selection, working memory, and executive control processes (McDougle et al., 2015; Taylor et al., 2014; Taylor & Ivry, 2011). The increased explicit contribution observed here, therefore, suggests that cognitive operations involved in identifying and selecting the correct target may recruit many of the same neural and computational mechanisms that support strategic compensation for visuomotor errors.

The ability of participants in the high-load condition to generate a larger explicit component despite significantly greater task difficulty suggests that strategic adaptation can be enhanced when cognitive processing remains aligned with task goals, offering a potential resolution to the discrepancy between the current findings and previous dual-task studies. Thus, task relevance can determine whether cognitive load impairs or facilitates explicit adaptation. Thus, the key factor may not be cognitive load per se, but whether the cognitive demands are congruent with the movement being adapted. Nevertheless, the interpretation of explicit and implicit contributions must be considered in light of the modeling approach employed. The decomposition of behavior into fast and slow processes assumes that these components correspond to explicit and implicit learning, respectively (McDougle et al., 2015). While this framework is widely used and is consistent with the results in the present study, future investigations incorporating direct measures of aiming strategies or verbal reports would provide a stronger test of the proposed increase in explicit learning.

### Cognitive Load and Steady State Learning

Participants in the high-load condition exhibited lower asymptotic adaptation despite showing a larger explicit contribution, indicating that enhanced strategic engagement does not necessarily lead to greater long-term compensation. This effect may be explained by classical theories of limited-capacity information processing (Kahneman, 1970). From this perspective, the additional cognitive demands of difficult visual search may reduce the resources available for learning. Thus, adaptation may stabilize at a lower level because the system has reached a point where it trades accuracy for ongoing cognitive demands. A second, and in our view more compelling, explanation derives from the interaction between explicit and implicit learning processes. The model fits revealed that greater explicit adaptation in the hard condition was accompanied by reduced implicit adaptation. Since a slower implicit process is characterized by greater retention across trials, a smaller contribution would be expected to lower the eventual adaptation asymptote. In contrast, the faster explicit process is generally more susceptible to forgetting. Consequently, an adaptation profile dominated by explicit processes may exhibit robust early compensation, reaching a lower steady-state level. This interpretation naturally explains why learning curves were broadly similar despite differences in asymptotic performance and why increased explicit learning failed to generate superior overall adaptation. These findings further suggest that the degree of adaptation observed behaviourally cannot be interpreted solely as an index of learning capacity. Instead, similar behavioral outcomes may arise from different combinations of underlying processes. Understanding how cognitive context alters the contribution of these processes may therefore be essential for interpreting variability in adaptation performance across different experimental paradigms.

### Relation to the competition model

The present findings provide additional support for the competition model proposed by Albert et al. (Albert, Jang, Modchalingam, ‘T Hart, et al., 2022; Benson et al., 2011; Neville & Cressman, 2018; Saijo & Gomi, 2010), which argues that explicit and implicit learning processes compete for a common error signal: increasing the contribution of one process reduces the error available to drive the other. In contrast, independent-error accounts propose that explicit adaptation is driven primarily by task errors, whereas implicit adaptation is driven by sensory prediction errors (Mazzoni & Krakauer, 2006; Morehead et al., 2017; Taylor et al., 2014). Several observations from the present study favor the competition account. First, enhanced explicit learning under high cognitive load was accompanied by a reduction in implicit learning. Second, the negative relationship observed between explicit and implicit learning across participants indicates that stronger expression of one process was associated with weaker expression of the other. Third, the changes in expression of these learning processes occurred despite similar behavioral adaptation curves, suggesting a redistribution of learning rather than a change in overall error exposure. Fourth, differences in peak and area under the curve between groups were evident not only in late adaptation but also in the estimated learning dynamics throughout the perturbation period. This suggests that competitive interactions emerge throughout learning, extending the model of Albert et al. (2022), which primarily focused on steady-state adaptation.

### Effect of Reaction Time and Action Errors

The expression of explicit processes is greater for movements associated with larger reaction times (Fernandez-Ruiz et al., 2011; Haith et al., 2015; Huberdeau et al., 2015), which is also a hallmark of a greater cognitive load condition (Figure 1C; right). Although not a classic load manipulation, preparation-time constraints could be considered another form of cognitive constraint (Figure 1C; left). Nonetheless, reaction times did not predict explicit adaptation at the individual participant level, suggesting that the prolonged movement initiation time alone could not account for the observed increase in explicit learning. Instead, the longer reaction times may reflect increased cognitive engagement, or the processing demands imposed by the high-load condition. In addition, the harder task also produced more action errors, leading to lower overall accuracy (Figure 1C; left). However, accuracy on the decision-making component of the secondary task did not affect adaptation, indicating that the interaction between the decision-making process and motor adaptation was not responsible for the observed differences across conditions. Taken together, our findings support the view that the cognitive state induced by the high-load condition, rather than the behavioral demands of the secondary task or the additional preparation time itself, is the critical factor modulating explicit learning.

### Relation to Executive Control

In this study, both tasks (Botvinick et al., 2001; Cohen et al., 2000) involve evaluative and executive components, suggesting evaluation within a unified cognitive control framework (Botvinick et al., 2001; Cohen et al., 2000). Numerous studies suggest that the Anterior Cingulate Cortex (ACC), a region involved in error processing (Gehring et al., 1993, 1995) and performance monitoring (Ito et al., 2003), may contribute to explicit motor learning, showing greater activity observed when larger errors are made (Seidler et al., 2013). ACC may integrate signals across tasks and recruit executive control under high load and conflict, enhancing prefrontal processes that update movement plans planned by sensorimotor networks. This could facilitate explicit, strategy-based adjustments while reducing reliance on implicit mechanisms, shifting the balance of learning toward explicit processes. More broadly, the results are compatible with the idea that explicit adaptation represents a form of cognitive control applied to motor behavior. From this perspective, one can understand why task-relevant cognitive demands enhanced explicit adaptation. When cognitive processing contributes directly to goal selection and action planning, greater engagement of control systems may improve the implementation of explicit corrective strategies. In contrast, when cognitive processing is directed toward an unrelated task, as in conventional dual-task paradigms, those same systems may be diverted away from adaptation.

In conclusion, the present study demonstrates that the influence of cognitive load on motor adaptation depends on the relationship between cognitive load and the motor task. Task-relevant cognitive load enhanced explicit learning while reducing implicit adaptation, a pattern that contrasts with traditional dual-task findings and highlights the importance of attentional and behavioural context. These results support the notion that explicit adaptation is closely linked to cognitive operations involved in action selection and executive control. Furthermore, the reciprocal changes in explicit and implicit learning provide additional evidence for a competition-based account of motor adaptation, in which both processes draw on a shared error signal. Collectively, the findings suggest that cognitive processes shape motor learning not only by determining how much adaptation occurs, but also by determining which learning system is primarily engaged to achieve behavioural goals.

## ACKNOWLEDGEMENTS

This work was supported by funding from a grant from the DBT-IISc partnership grant, an ANRF grant, and intramural support from IISc.

## AUTHOR CONTRIBUTIONS

V.R., S.S., and A.M. conceptualised and designed the study. V.R performed the experiments and collected the data. V.R analysed the data. V.R. drafted the manuscript, and S.S. and A.M. critically reviewed it for important intellectual content. A.M and S.S supervised the project. All authors reviewed, edited, and approved the final version of the manuscript.

## DATA AVAILABILITY STATEMENT

All data and codes will be publicly available upon manuscript acceptance.

## REFERENCES

Albert, S. T., Jang, J., Modchalingam, S., ‘t Hart, B. M., Henriques, D., Lerner, G., Della-Maggiore, V., Haith, A. M., Krakauer, J. W., & Shadmehr, R. (2022). Competition between parallel sensorimotor learning systems. eLife, 11, e65361. 10.7554/eLife.65361

Botvinick, M. M., Braver, T. S., Barch, D. M., Carter, C. S., & Cohen, J. D. (2001). Conflict monitoring and cognitive control. Psychological Review, 108(3), 624–652. 10.1037/0033-295X.108.3.624

Cohen, J. D., Botvinick, M., & Carter, C. S. (2000). Anterior cingulate and prefrontal cortex: Who’s in control? Nature Neuroscience, 3(5), 421–423. 10.1038/74783

Gehring, W. J., Coles, M. G., Meyer, D. E., & Donchin, E. (1995). A brain potential manifestation of error-related processing. Electroencephalography and Clinical Neurophysiology. Supplement, 44, 261–272.

Gehring, W. J., Goss, B., Coles, M. G. H., Meyer, D. E., & Donchin, E. (1993). A Neural System for Error Detection and Compensation. Psychological Science, 4(6), 385–390. 10.1111/j.1467-9280.1993.tb00586.x

Haith, A. M., Huberdeau, D. M., & Krakauer, J. W. (2015). The Influence of Movement Preparation Time on the Expression of Visuomotor Learning and Savings. The Journal of Neuroscience, 35(13), 5109–5117. 10.1523/JNEUROSCI.3869-14.2015

Huberdeau, D. M., Krakauer, J. W., & Haith, A. M. (2015). Dual-process decomposition in human sensorimotor adaptation. Current Opinion in Neurobiology, 33, 71–77. 10.1016/j.conb.2015.03.003

Ito, S., Stuphorn, V., Brown, J. W., & Schall, J. D. (2003). Performance Monitoring by the Anterior Cingulate Cortex During Saccade Countermanding. Science, 302(5642), 120–122. 10.1126/science.1087847

Kahneman, D. (1970). Remarks on attention control. Acta Psychologica, 33, 118–131. 10.1016/0001-6918(70)90127-7

Keisler, A., & Shadmehr, R. (2010). A Shared Resource between Declarative Memory and Motor Memory. The Journal of Neuroscience, 30(44), 14817–14823. 10.1523/JNEUROSCI.4160-10.2010

Kim, H. E., Morehead, J. R., Parvin, D. E., Moazzezi, R., & Ivry, R. B. (2018). Invariant errors reveal limitations in motor correction rather than constraints on error sensitivity. Communications Biology, 1(1), 19. 10.1038/s42003-018-0021-y

Leow, L.-A., Marinovic, W., De Rugy, A., & Carroll, T. J. (2020). Task Errors Drive Memories That Improve Sensorimotor Adaptation. The Journal of Neuroscience, 40(15), 3075–3088. 10.1523/JNEUROSCI.1506-19.2020

Mazzoni, P., & Krakauer, J. W. (2006). An Implicit Plan Overrides an Explicit Strategy during Visuomotor Adaptation. Journal of Neuroscience, 26(14), 3642–3645. 10.1523/JNEUROSCI.5317-05.2006

McDougle, S. D., Bond, K. M., & Taylor, J. A. (2015). Explicit and Implicit Processes Constitute the Fast and Slow Processes of Sensorimotor Learning. Journal of Neuroscience, 35(26), 9568–9579. 10.1523/JNEUROSCI.5061-14.2015

Morehead, J. R., Taylor, J. A., Parvin, D. E., & Ivry, R. B. (2017). Characteristics of Implicit Sensorimotor Adaptation Revealed by Task-irrelevant Clamped Feedback. Journal of Cognitive Neuroscience, 29(6), 1061–1074. 10.1162/jocn_a_01108

Neville, K.-M., & Cressman, E. K. (2018). The influence of awareness on explicit and implicit contributions to visuomotor adaptation over time. Experimental Brain Research, 236(7), 2047–2059. 10.1007/s00221-018-5282-7

Oldfield, R. C. (1971). The assessment and analysis of handedness: The Edinburgh inventory. Neuropsychologia, 9(1), 97–113. 10.1016/0028-3932(71)90067-4

Sadaphal, D. P., Kumar, A., & Mutha, P. K. (2022). Sensorimotor Learning in Response to Errors in Task Performance. Eneuro, 9(2), ENEURO.0371-21.2022. 10.1523/ENEURO.0371-21.2022

Saijo, N., & Gomi, H. (2010). Multiple Motor Learning Strategies in Visuomotor Rotation. PLoS ONE, 5(2), e9399. 10.1371/journal.pone.0009399

Seidler, R. D., Kwak, Y., Fling, B. W., & Bernard, J. A. (2013). Neurocognitive Mechanisms of Error-Based Motor Learning. In M. J. Richardson, M. A. Riley, & K. Shockley (Eds), Progress in Motor Control (Vol. 782, pp. 39–60). Springer New York. 10.1007/978-1-4614-5465-6_3

Shingane, S. N., Rao, N., Kumar, N., & Mutha, P. K. (2025). Task relevance selectively modulates sensorimotor adaptation in the presence of multiple prediction errors. Journal of Neurophysiology, 134(5), 1607–1618. 10.1152/jn.00511.2024

Smith, M. A., Ghazizadeh, A., & Shadmehr, R. (2006). Interacting Adaptive Processes with Different Timescales Underlie Short-Term Motor Learning. PLoS Biology, 4(6), e179. 10.1371/journal.pbio.0040179

Taylor, J. A., & Ivry, R. B. (2011). Flexible Cognitive Strategies during Motor Learning. PLoS Computational Biology, 7(3), e1001096. 10.1371/journal.pcbi.1001096

Taylor, J. A., Krakauer, J. W., & Ivry, R. B. (2014). Explicit and Implicit Contributions to Learning in a Sensorimotor Adaptation Task. The Journal of Neuroscience, 34(8), 3023–3032. 10.1523/JNEUROSCI.3619-13.2014

Taylor, J. A., & Thoroughman, K. A. (2007). Divided Attention Impairs Human Motor Adaptation But Not Feedback Control. Journal of Neurophysiology, 98(1), 317–326. 10.1152/jn.01070.2006

Zhang, X., Zhang, T., & Wei, K. (2025). Cognitive load suppresses explicit learning while sparing implicit learning in visuomotor adaptation. Journal of Neurophysiology, 134(4), 1133–1145. 10.1152/jn.00262.2025

